# *Aedes aegypti* Aag-2 culture cells enter endoreplication process upon pathogen challenge

**DOI:** 10.1101/2021.01.13.425146

**Authors:** Christian Domínguez-Benítez, Javier Serrato-Salas, Renaud Condé, Humberto Lanz-Mendoza

**Affiliations:** Laboratorio de Infección e Inmunidad. Centro de Investigaciones sobre Enfermedades Infecciosas. Instituto Nacional de Salud Pública, Av. Universidad 655, CP 62100. Cuernavaca, Morelos, México

## Abstract

Metamorphic insects apparently rely on a finite number of cells after emergence to counterbalance either commensal and pathogen presence. For hematophagous insects, blood-feeding is a crucial step for offspring development, therefore enteric cells repairing molecular mechanisms consists in fine regulated pathways to counterattack biotic and abiotic insults. Nevertheless, recent research suggests that midgut cells are capable to adapt their immune responses to pathogen challenges. Recently, *Anopheles* and *Aedes* mosquitoes have been observed to increase their DNA cell content upon encounter with parasites, bacteria and virus respectively. Genomic endoreplication is one of the most important processes in larval development for fast transcriptional activity and protein secretion.

So, in this paper we explore the ability of *Aedes aegypti* Aag-2 culture cells to develop a likely endoreplication process to face pathogen presence. Aag-2 cells at 6 and 12 hours post-biotic insult enter a proliferation arrest and increases DNA content, these two phenomena recovers control levels at 24 h post-treatment. It requires more research data about the type of genomic regions that has been replicated in the process, and the concentration that antimicrobial molecules are released into culture media.

## Introduction

Host biological responses against invading pathogens relies on inducible and constitutive responses, humoral and cellular components, divided into innate and adaptive responses (Sprent, 1994). Typical innate peripheral cells are NK and monocyte/macrophages and dendritic cells (Weavers, Evans, Martin, & Wood, 2016); humoral defenses comprise antibody production and complement system, besides a large families of molecules with antimicrobial properties (Burnet, 1976). In invertebrates, host responses against invading pathogens relies on limited resources through innate responses like phagocytosis, nodule formation, encapsulation, reactive oxygen species local responses and antimicrobial peptides production (Barnard, Nijhof, Fick, Stutzer, & Maritz-Olivier, 2012; Costa, Jan, Sarnow, & Schneider, 2009; De Gregorio, Spellman, Tzou, Rubin, & Lemaitre, 2002; Dong et al., 2006; Hillyer, 2010; Kleino, 2010). These immune responses are controlled mainly by four transcriptional cascades, Toll, IMD, Jak/STAT and RNAi (Carissimo et al., 2015; Fragkoudis, Attarzadeh-Yazdi, Nash, Fazakerley, & Kohl, 2009; Lee, Lee, & Lee, 2017; Pasquier, 2005; Schmid-Hempel, 2005; Vilcinskas, 2013; Waldock, Olson, & Christophides, 2012).

One of the most promising elements to face invading pathogens is the ability to unpack DNA to transcribe RNAm and translate a higher protein output to neutralize invading pathogen population (van Zelm, Szczepański, van der Burg, & van Dongen, 2007). In mammals, adaptive responses relies on immune cells entering clonal expansion and antibody affinity selection to cope with the proliferating pathogen population (Bassing, Swat, & Alt, 2002; Sprent, 1994; van Zelm et al., 2007; Weng, Araki, & Subedi, 2012).

In *Drosophila melanogaster* (*Dmel*) the DNA synthesis has been studied in detail during larval and embryonic stages, where mitotic, incomplete mitotic or endoreplication cycles fluctuate in egg chamber in fine regulated mechanisms to developing wings, dorsal polarization and complete egg shell formation (Jia, Tamori, Pyrowolakis, & Deng, 2014; Perdigoto & Bardin, 2013; Shen & Sun, 2017). Recent research suggests that enteric cells and circulating hemocytes are not only provided by stem cell differentiation during earlier stages, but presence of adult cell clusters progenitor post-larval hematopoiesis relies on Delta-Notch signaling in a similar fashion as in pupal stem cells, and are capable to respond to pathogen challenges (Ghosh, Singh, Mandal, & Mandal, 2015; Guo & Ohlstein, 2015).

In other dipterid, such as *Aedes aegypti*, the midgut adult cells are supposed to not enter mitosis, hence the adaptive-like response observed to recurrent pathogen challenge would imply a genomic change in the challenged cells that would allow the production of effector molecules at a faster pace than the one developed during an only first encounter. We have previously demonstrated that, upon viral challenge and inactive virus oral fed, the mosquito *Aedes aegypti* midgut and carcass cells entered an endoreplication cycle (Serrato-Salas, Hernández-martínez, et al., 2018; Serrato-Salas, Izquierdo-Sánchez, et al., 2018).

This process is controlled by *hnt* gene through Delta-Notch signaling cascade, triggering the genomic DNA synthesis without entering in a mitotic process. The process limits the viral spread to the neighboring cells and is probably related to the progressive diminution of the viral load in the intestinal and somatic cell after the initial infection (Salazar, Richardson, Sánchez-Vargas, Olson, & Beaty, 2007).

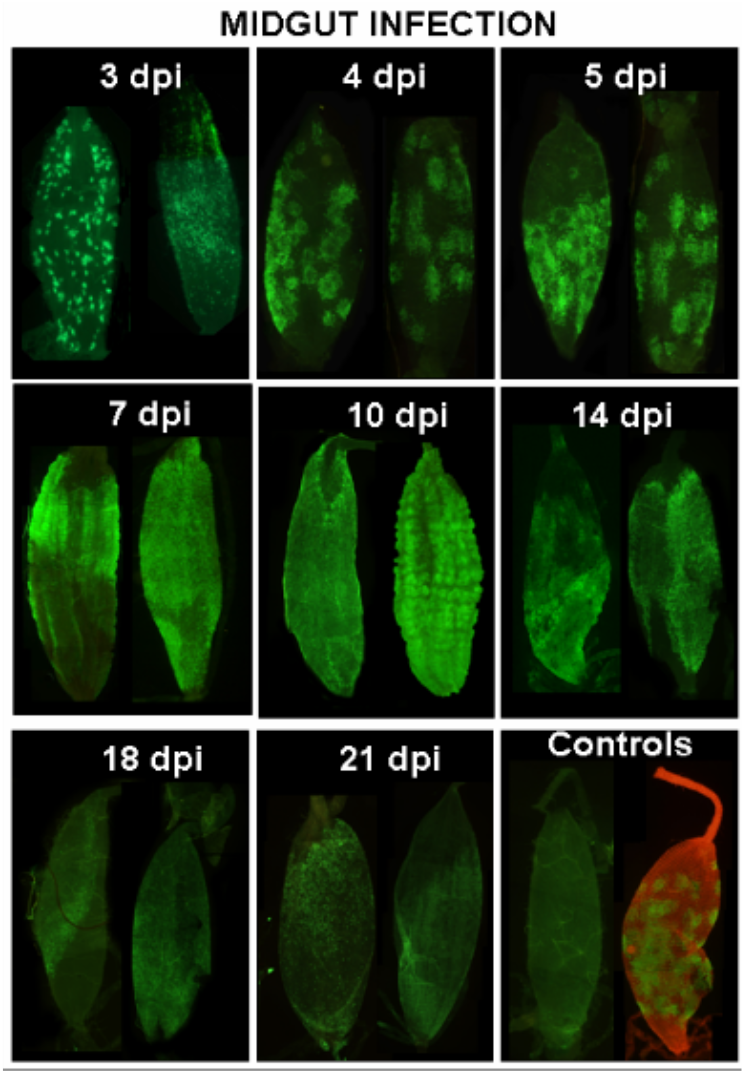

Here we studied the effect of a pathogenic challenge on *Aedes aegypti* culture cells proliferation and immune response.

## Materials and methods

### Aag-2 cell culture and fungal challenge

*Aedes aegypti* Aag-2 cells (ATCC CCL-125) were cultured at 28°C in Schneider’s *Drosophila* medium for maintenance and microbial challenge. The cell media were supplemented with 10% heat-inactivated fetal bovine serum, 2 mM L-glutamine, and 100 U/mL each of penicillin, streptomycin and neomycin. The monolayer Aag-2 cells were maintained by re-seeding two-three times per week. The cells plates confluence used for assays was minimal 80%. Zymosan-PBS solution was added into the cultured cell medium for 1 h in 24 wells plates.

A proliferation assay was performed for Aag-2 cells treated with different zymosan concentrations at 24 h. The 50% maximum cytotoxic concentration was determined, to test the effects in cells were not caused by direct reduced proliferation.

After incubation, zymosan was removed and replaced with fresh Schneider’s media. The stimulated cells were collected through Trypsin-Versene (Thermo Scientific) partial digestion.

### Cells viability assay

Cell viability was determined by trypan blue dye exclusion (0.4% in PBS pH 7.5) in a Neubauer chamber using a bright field microscope. Three-hundred cells were counted for each treatment from five independent experiments done by duplicate. MTT [4,5 dimethylthiazolil-2]-2,5-diphenyl tetrazolium bromide reduction assays were performed in 96 wells plates. After stimulus, media was removed and replaced with fresh media plus MTT stock solution. After 3 h incubation at 28°C, cells reduced MTT to a formazan insoluble product. Supernatants were transferred to an ELISA plate and solubilized in SDS 10%-HCl 0.01 N solution mix. Optical density was recorded at a 595nm wavelength. At least, five biological assays were performed. Previously, a standard curve calibration was performed to correlate optical density to number of cells.

### cDNA synthesis and qPCR strategy

Total RNA from collected Aag-2 cells were performed following protocol instructions from Trizol Reagent (Sigma-Aldrich). Complementary DNA (cDNA) was synthesized for qPCR detection. Quantitative PCR were performed in a ViiA 7 Real-Time PCR System (Applied Biosystems) with the QuantStudio Real Time Software v1.3. Reactions were realized in a total volume of 10μL containing 5μL of SYBR Green PCR Master mix, 1.5 ng of cDNA template, 250 nmol of each one of primers, and volume completed with nuclease-free water. Reaction conditions were the following: 50°C for 2 min, 95°C for 10 min followed by 40 cycles of denaturation at 95°C for 15 s, annealing and extension at 60°C for 1 min. The Ct values obtained from the tested gene relative to the reference gene, was used to obtain delta Ct values of zymosan treated and control cell samples. S7 gene (ribosomal unit S70) was selected as the reference gene (Moreno-García, Vargas, Ramírez-Bello, Hernández-Martínez, & Lanz-Mendoza, 2015; Vargas, Moreno-García, Duarte-Elguea, & Lanz-Mendoza, 2016). Relative expression values were obtained using the delta-delta cycle threshold (DDCT) method (Bubner, Gase, & Baldwin, 2004).

We used the PCR primers described in our previous work to quantify the transcription of *hnt* gene upon microbial molecule challenge (Serrato-Salas, Hernández-martínez, et al., 2018; Serrato-Salas, Izquierdo-Sánchez, et al., 2018). Briefly, Ribosomal protein S7 (AAEL009496) 292 bp amplicon; forward 5’ GGG ACA AAT CGG CCA GGC TAT C 3’, reverse 5’ TCG TGG ACG CTT CTG CTT GTT G 3’ primers were used for internal control PCR (Xi, Ramirez, & Dimopoulos, 2008). From CDS and mRNA predicted sequences, following primers were designed: hnt (Fwd): 5’ CGC AAG GAG TTA GAG CGT GA 3’, hnt (Rev): 5’ GTG TCG ATC GCA GTT GGA CT 3’.

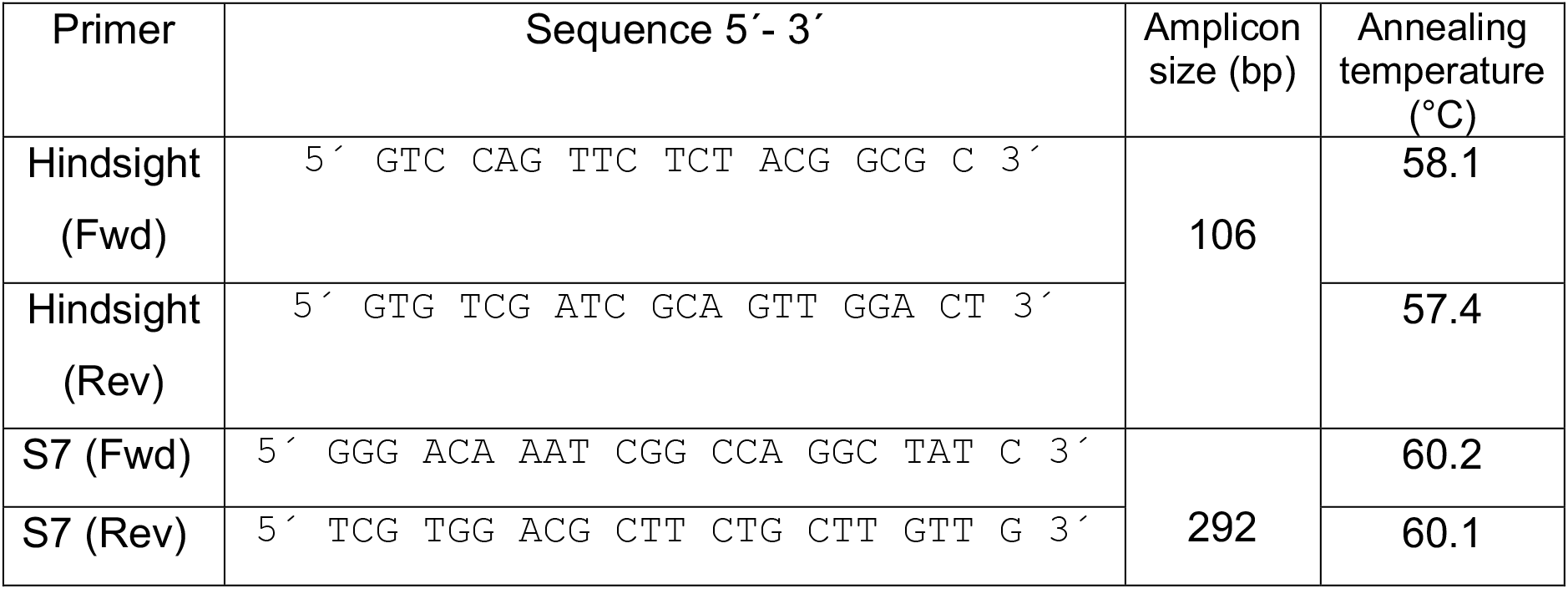

### Genomic DNA extraction

Total genomic DNA (gDNA) was extracted with Phenol-Chloroform-Isoamyl Alcohol mixture according to manufacturer’s instructions. Briefly, the culture cells pools placed in the solvent mixture were homogenized using a small piston until macerated. The phenol interface was washed with citrate buffer. The experimental gDNAs were precipitated with ethanol as per protocol (Hernandez-Martinez et al., 2006; Hernández-Martínez, Barradas-Bautista, & Rodríguez, 2013).

### gDNA fragmentation

The extracted gDNA was subsequently sonicated in a X sonicator, through five 30 seconds cycles (on/off) at low power at 4°C. The gDNA fragmentation was asserted in a 1% agarose gel run in TBE Buffer (Supplementary Material S1).

### gDNA melting curve

The fragmented gDNA melting curves were performed on a Real Time PCR CFX96 from Bio-Rad. 5μL of gDNA were mixed with 1μL of Taq buffer and 4μL of SYBR Green PCR Master mix. The mix was subjected to a high-resolution melting analysis from 65°C to 95°C at 0.3°c per second.

### Agarose pulsed-field electrophoresis

Intact cells were embedded in low melting agarose blocks (Schwartz & Cantor 1984). Cells are lysed and proteins are removed by Proteinase K treatment. This procedure yields intact DNA. The agarose block can be loaded directly into the well of a pulsed-field gel. Gels were cast using 1% SeaKem GTG agarose and the electrophoresis performed 24 hours at 120V in 0.5 x TBE buffer. After the completion of the run the agarose gel was stained for 15 min in ethidium bromide (1 mg/ml H2O). The gel was destained by two washes in 0.5xTBE for 1 hr with gentle agitation and photographed using a shortwave UV-light (254 nm). When the DNA was intended to be used for subsequent restriction and cloning, longwave UV-light (360 nm) was used to avoid nicking of the DNA.

## Results

### Dose-cytotoxic effect for zymosan-treated cells

A proliferation assay for Aag-2 cells treated with different zymosan concentrations for 24 h. The half maximum cytotoxic concentration (CC50) value was 34.07-36.91 g/L. (Figure 1A). The final zymosan concentration was 0.5 g/L (70 times lower), confirming that cell effects were not caused by reduced proliferation.

**Fig. 1:**
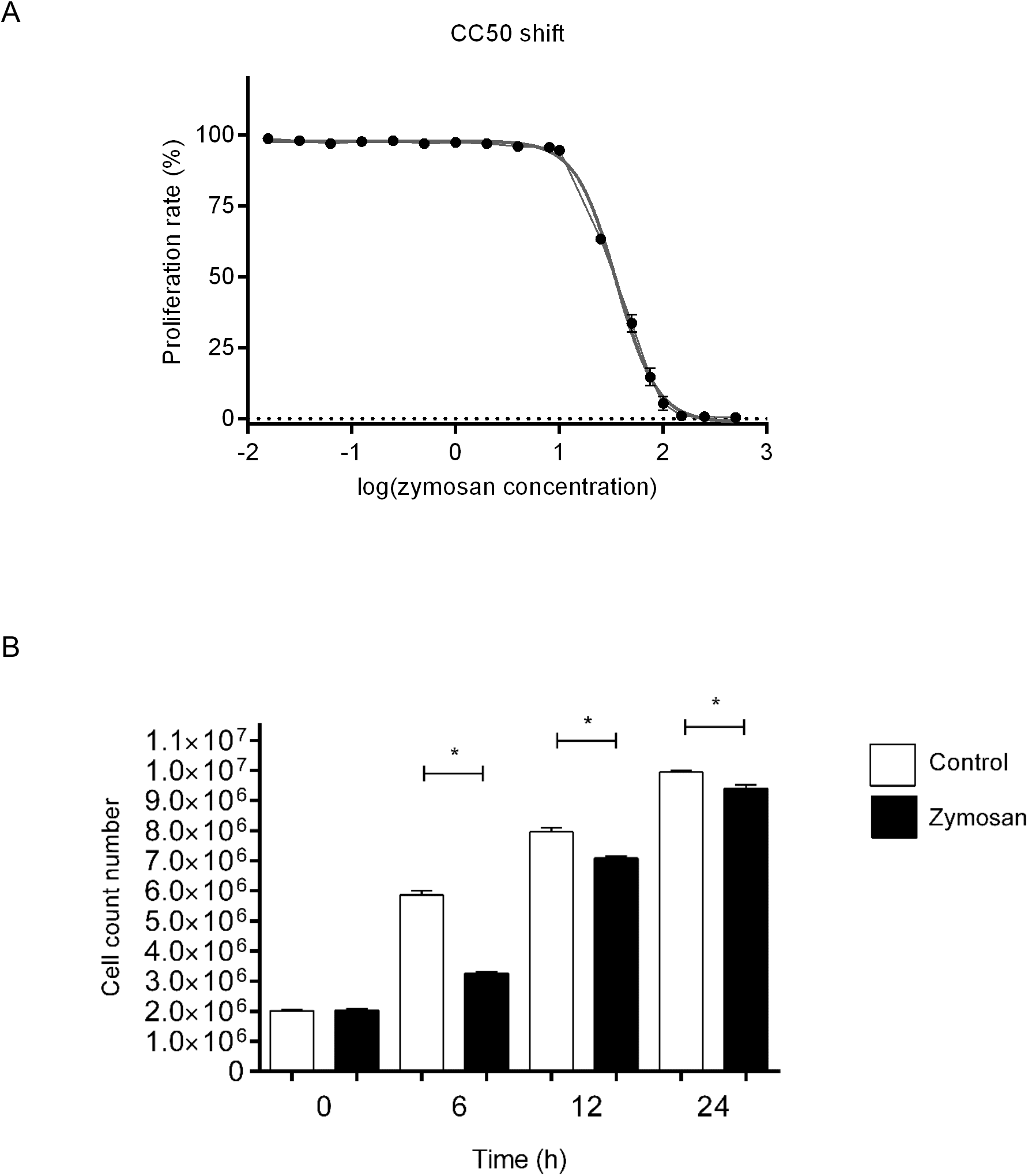
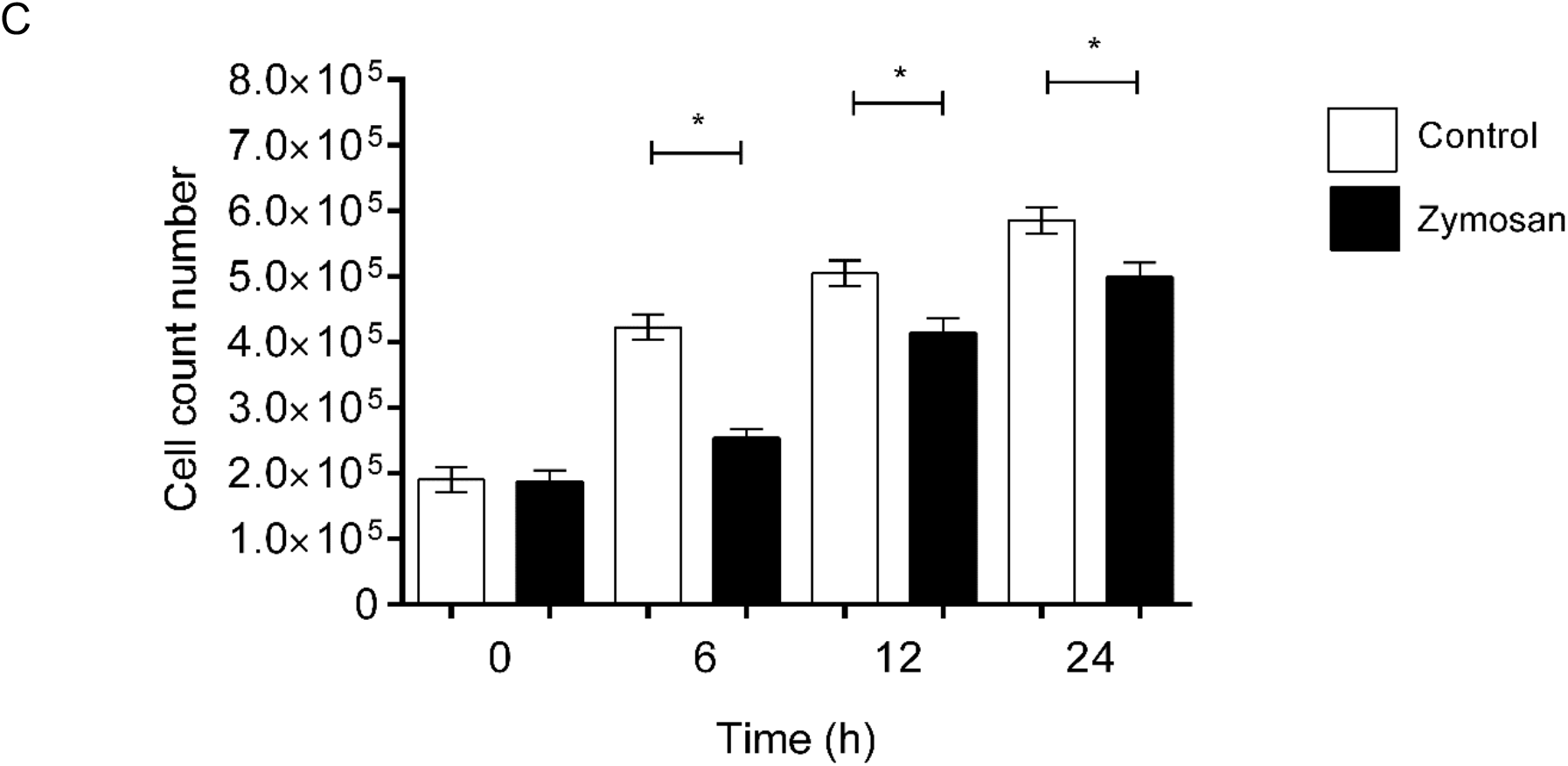
A) Cytotoxic concentration assay. LogCC50 1.532-1.567 (CC50 34.07-36.91 g/L). R^2^ = 0.9977. B) Aag-2 cells number per culture plate over 24 hours post zymosan challenge. Student’s T-test with Welch’s correction 0 h p=0.7148. 6 h p<0.0001. 12 h p<0.0001. 24 h p<0.0001. C) MTT exclusion dye viable cell number per culture plate over 24 hours post zymosan challenge. Student T-test welch correction values are for 0 h time point p=0.6694; 6 h p<0.0001; 12 h p<0.0001, 24 h p<0.0001.

### Zymosan treatment detains the Aag-2 cells proliferation

In order to assess the impact of the signaling mediated by fungal cell wall lysate, we treated Aag-2 cells with zymosan. We counted the cells post challenge and their viability were assessed. The difference in the number of cells counted post treatment points to a stoppage of cells duplication (Figure 1B). These cells were viable, and the differences of cell number remained equal when counting with MTT assay, hence discarding that the effect would be due to cell death (Figure 1C). Both tests showed that the cell proliferation was detained by the zymosan challenge and that 24 hours after, this difference was bridged. It seems that immune stress does affect cells duplication rate.

### Zymosan-mediated cell growth arrest changes the genomic DNA’s cell content

As described in Material and Methods, genomic DNA was obtained through organic solvent extraction (Phenol-Chloroform-Isoamyl alcohol mixture). The total gDNA amount was measured spectrophotometrically (wavelength 260/280 nm) and visualized in agarose gel electrophoresis through EtBr staining. Interestingly, though the number of viable cells was diminished upon zymosan treatment, the total DNA amount per plate increased. The amount of DNA per cell increased, the most prominent difference happening 6 h post challenge, remained at 12 h and recovers control levels at 24 h (Figure 2A). The highest gDNA relative ratio content per cell was observed 6 hours post treatment (Figure 2B).

**Figure 2:**
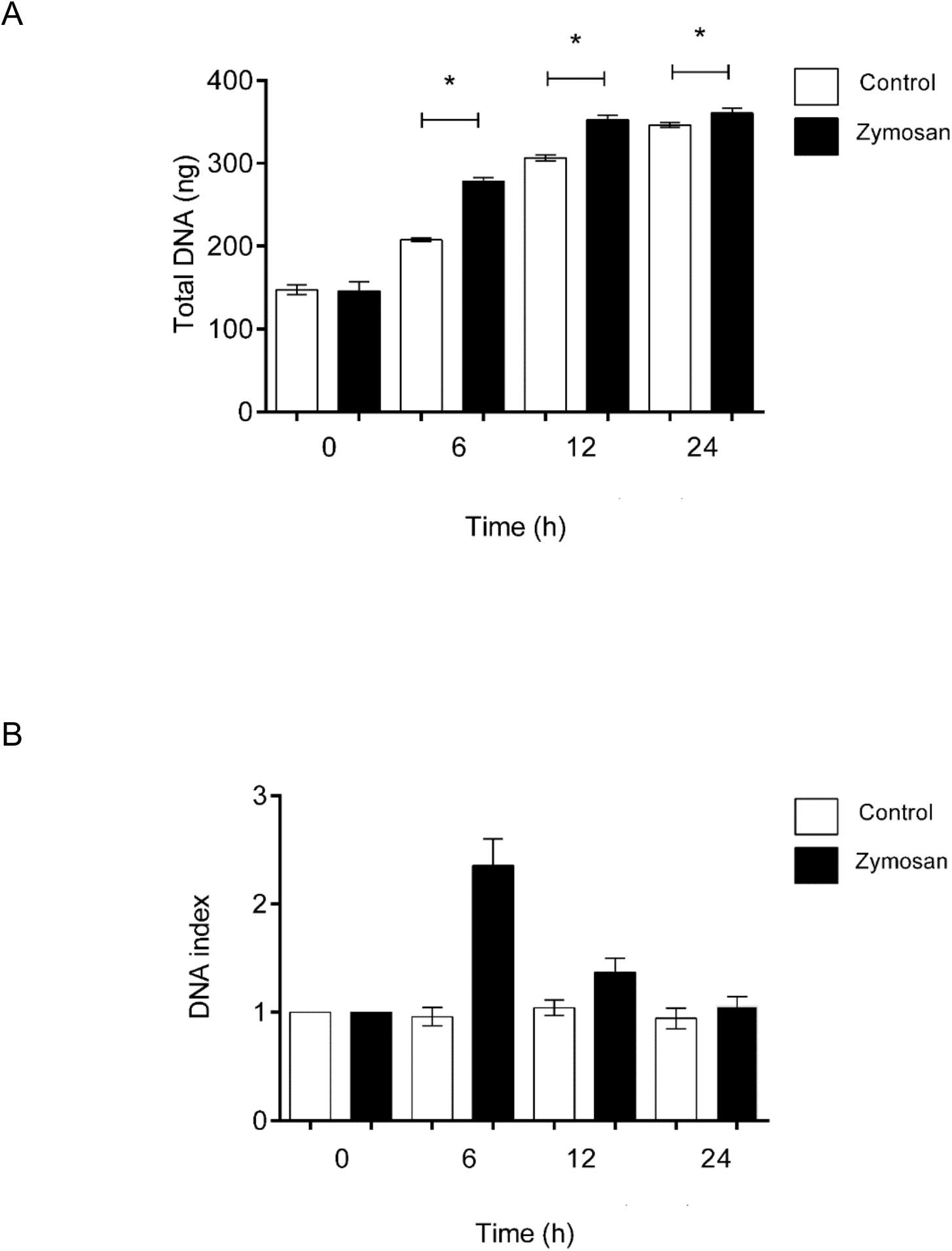
A) Total gDNA amount extracted from the cell cultures at 0, 6, 12 and 24 hours post zymosan challenge. 0 h p=0.6676. 6 h p<0.0001. 12 h p<0.0001. 24 h p<0.0001. B) Normalized relative gDNA content per cell at 0, 6, 12 and 24 hours post zymosan challenge.

### Zymosan challenge induces *Hnt* gene transcription

Endoreplication phenomena in the *Aedes aegypti* mosquito has been previously observed by our group (Hernandez-Martinez et al., 2006; Hernández-Martínez et al., 2013; Serrato-Salas, Hernández-martínez, et al., 2018; Serrato-Salas, Izquierdo-Sánchez, et al., 2018). We determined that the *hnt* gene transcription correlated with DNA base incorporation in the mosquito cells nuclei (Hernandez-Martinez et al., 2006; Serrato-Salas, Hernández-martínez, et al., 2018). Therefore, we analyzed the *hnt* gene transcription upon microbial challenge. We found that six hours after the zymosan challenge, *hnt* transcript nearly triplicated (Figure 3). Interestingly, the zymosan challenge elicited *hnt* transcription in a timely fashion as to be related with the gDNA/cell increase observed earlier.

**Figure 3:**
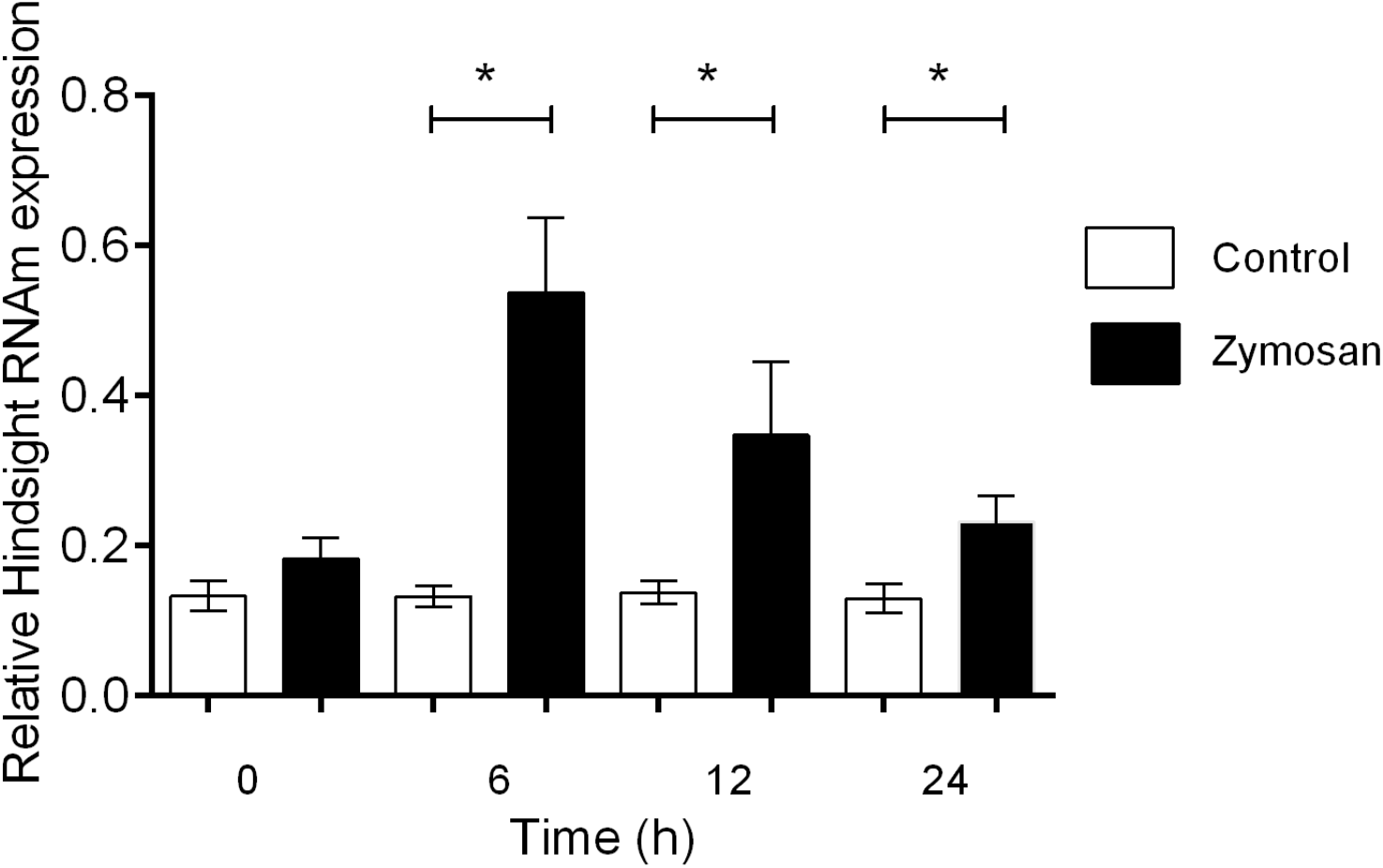
Relative *Hindsight* gene transcription post challenge. The transcripts amount values were normalized against ribosomal S7 gen. 0 h p=0.0001. 6 h p<0.0001. 12 h p<0.0001. 24 h p<0.0001.

### Zymosan stimulation altered gDNA sequence proportions

Cell genomic DNA entered an endoreplicating cycle with specific sequences amplified. The melting curve of the extracted gDNA showed diverging pattern in control and zymosan challenged Aag-2 culture cells. If the phenomenon would have implied a gDNA duplication, the proportion of each genomic sequences would have been equal hence their melting curves also. As can be observed in Figure 4, The melting curve of the sonicated gDNA of control and induced cells are different. The fact that some lower temperature peaks are bigger in zymosan treated cells point to an increase in the number of specific sequences against the rest of the genome, suggesting that only specific part of the genome are amplified. The melt temperature of the control samples was 75.06 +-2.1 °c while the zymosan treated gDNA had a melt temperature of 66.52+-3.2 °c. This could be due to the relative amount of repeated sequences of the endoreplicated sequences that would separate jointly at a lower temperature than the non endoreplicated gDNA part.

**Fig. 4.**
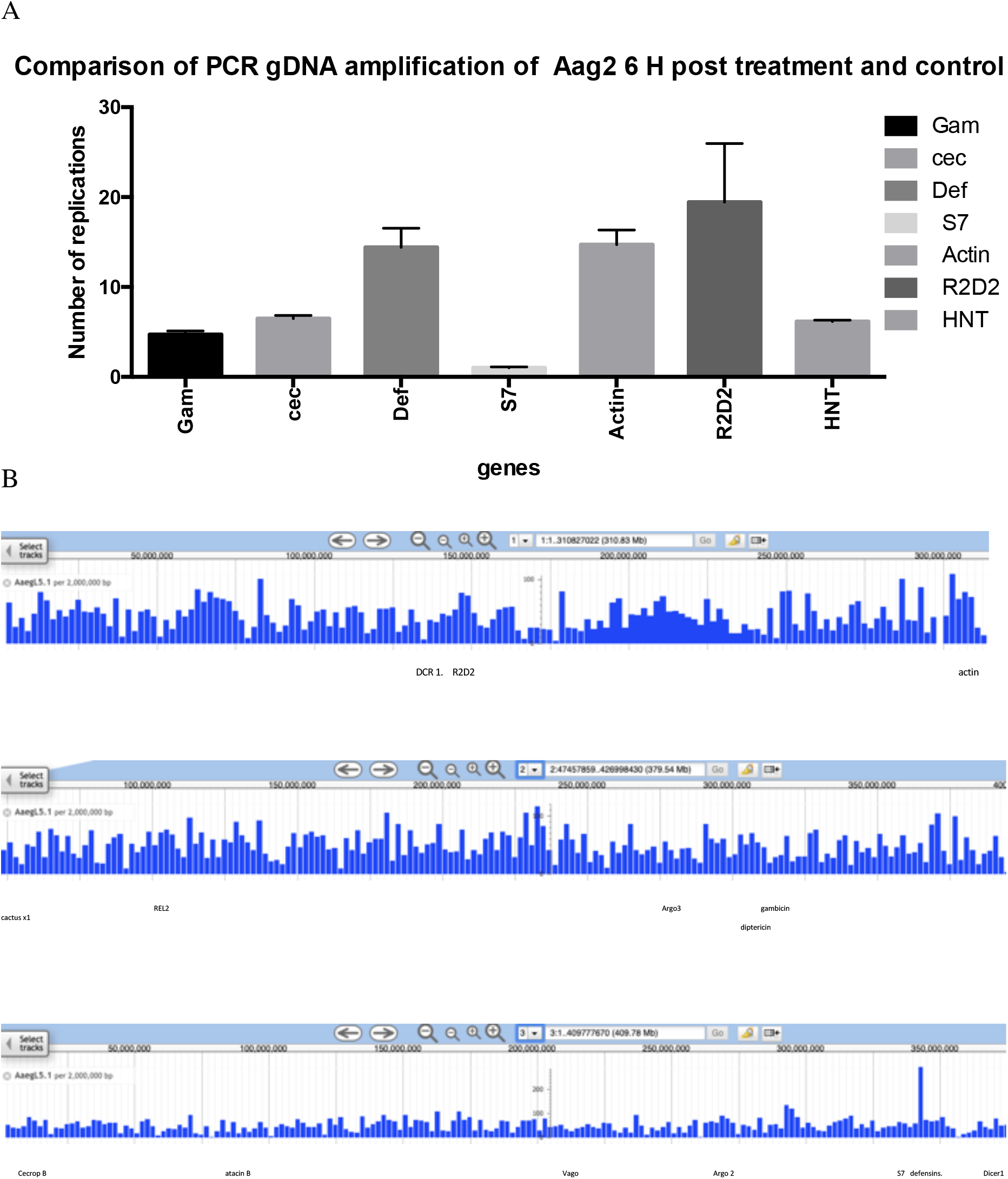
A Genomic PCR of distinct *Aedes* genes. The amplification of the same amount of gDNA was relativized to the gene that needed the most cycle to appear: S7. 4B Genomic distribution of the genes sequence chosen for gDNA qPCR.

**Figure 4.**
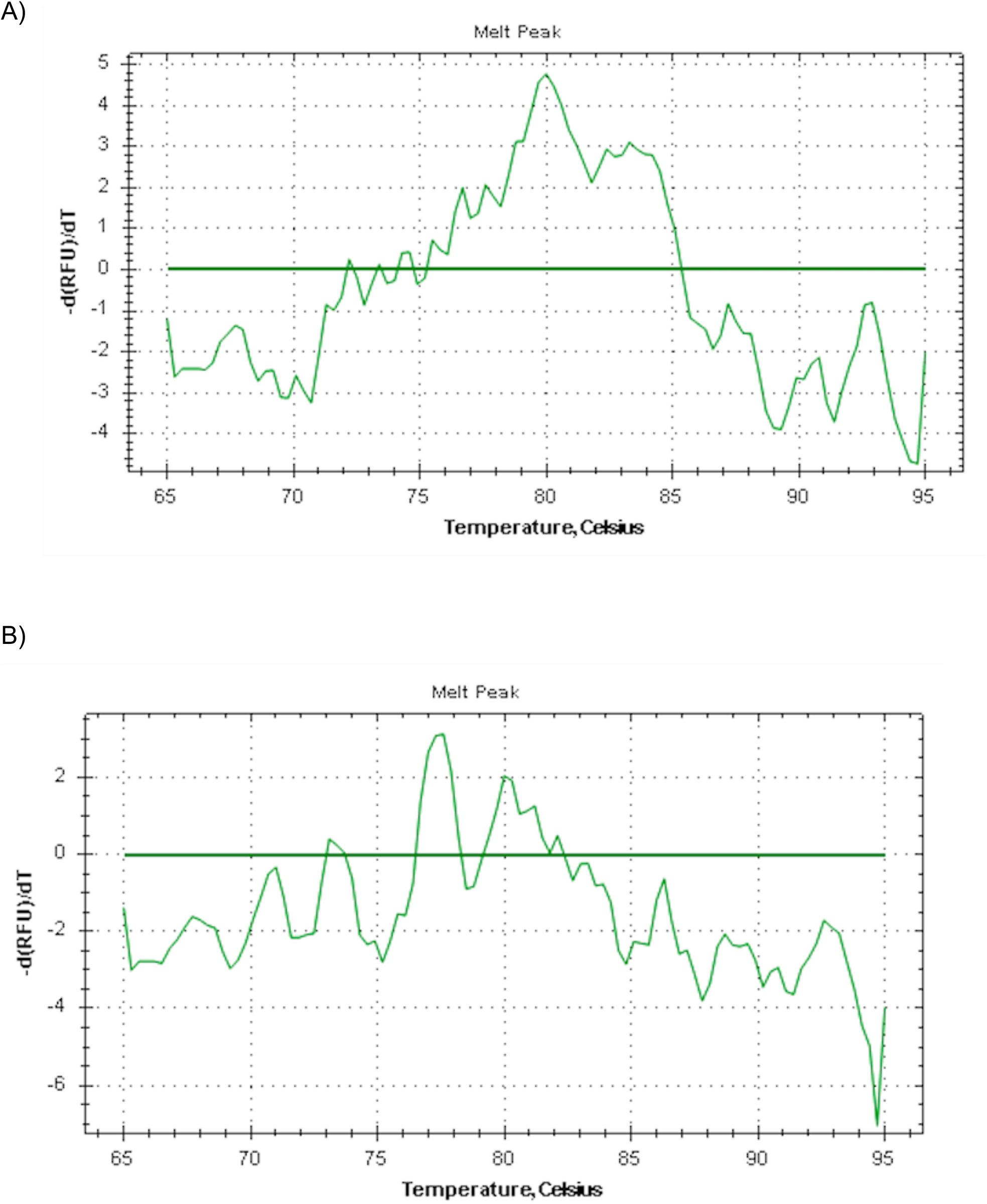
Melt peaks of sonicated cultured cells gDNA A) in control conditions B) six hours post Zymosan exposure.

### Zymosan stimulated *Aag2* cells gDNA present lower molecular weight DNA

The later evidences pointed to the amplification of specific gDNA regions by the *Aag2* cells submited to the zymosan stress. The partial gDNA amplification implied the presence of large extrachromosomal DNA sequences in the treated cells nucleus. Therefore, we separate the chromosomal DNA by pulse field chromatography, expecting to see large DNA fragment in the gDNA of the treated cells. As can be seen in figure 5, we did observe such fragment in the treated cell gDNA between 25 and 35 kBases.

**Figure 5:**
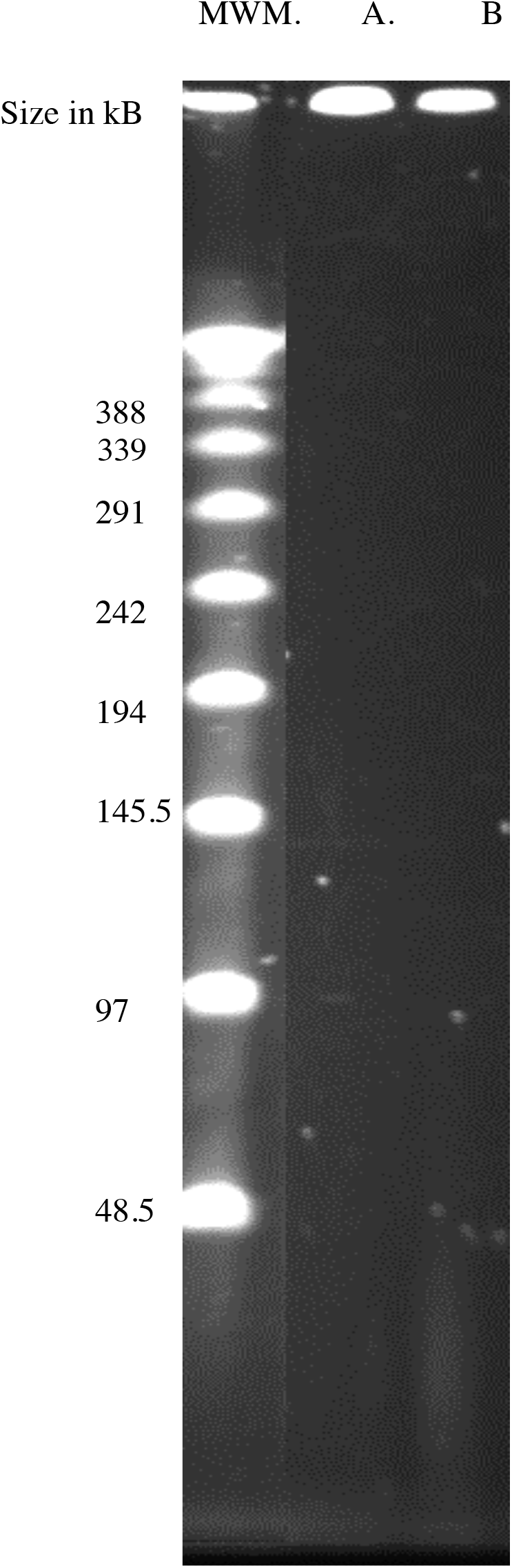
Pulsed field gel of the *Aag2* cells gDNA Control (A) and 6 hours post Zymozan challenge (B).

**Fig. 6:**
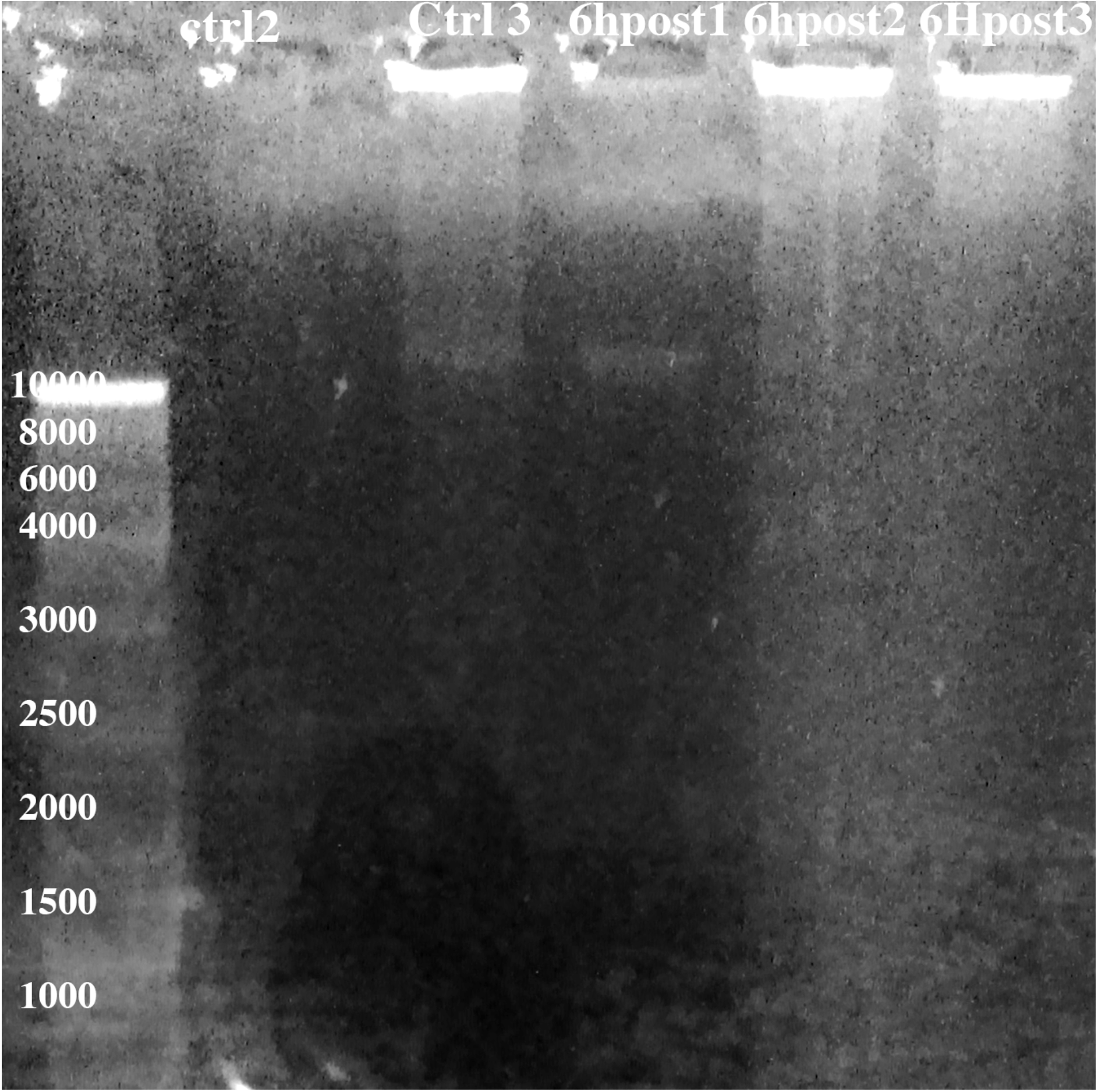
gDNA of control and zymosan treated cells.

## Conclusion

Aag-2 cells is a model for *Aedes aegpyti* competent immune response. Immune challenge induces cell division arrest in these cells. The relative genomic DNA content transiently increases at the same time that *hnt* gene transcription increases. The DNA content increase is related to specific genomic sequence amplification, the same are transient and resolved 24 hours post immune stress.

---

**Figure S1:**
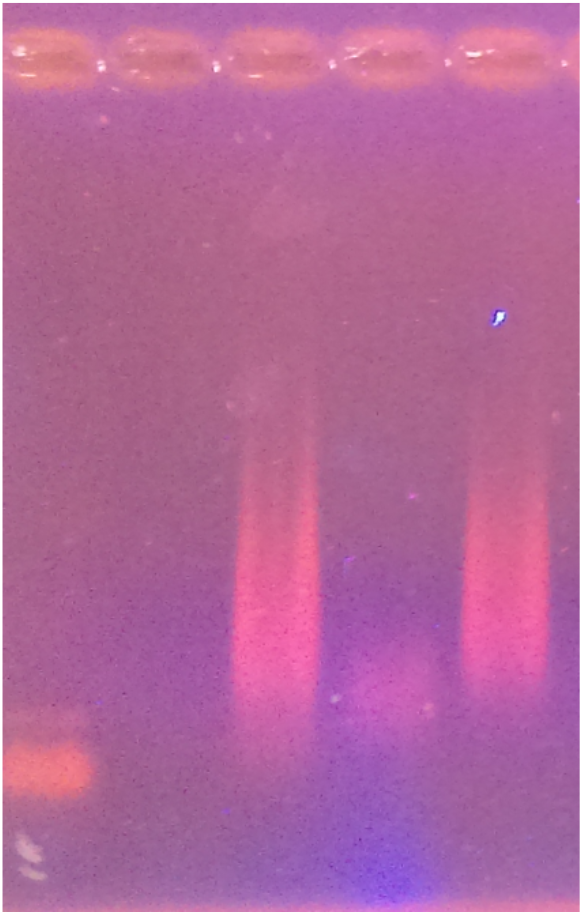
Aag-2 cells gDNA fragmentation.

**Fig.S.2:**
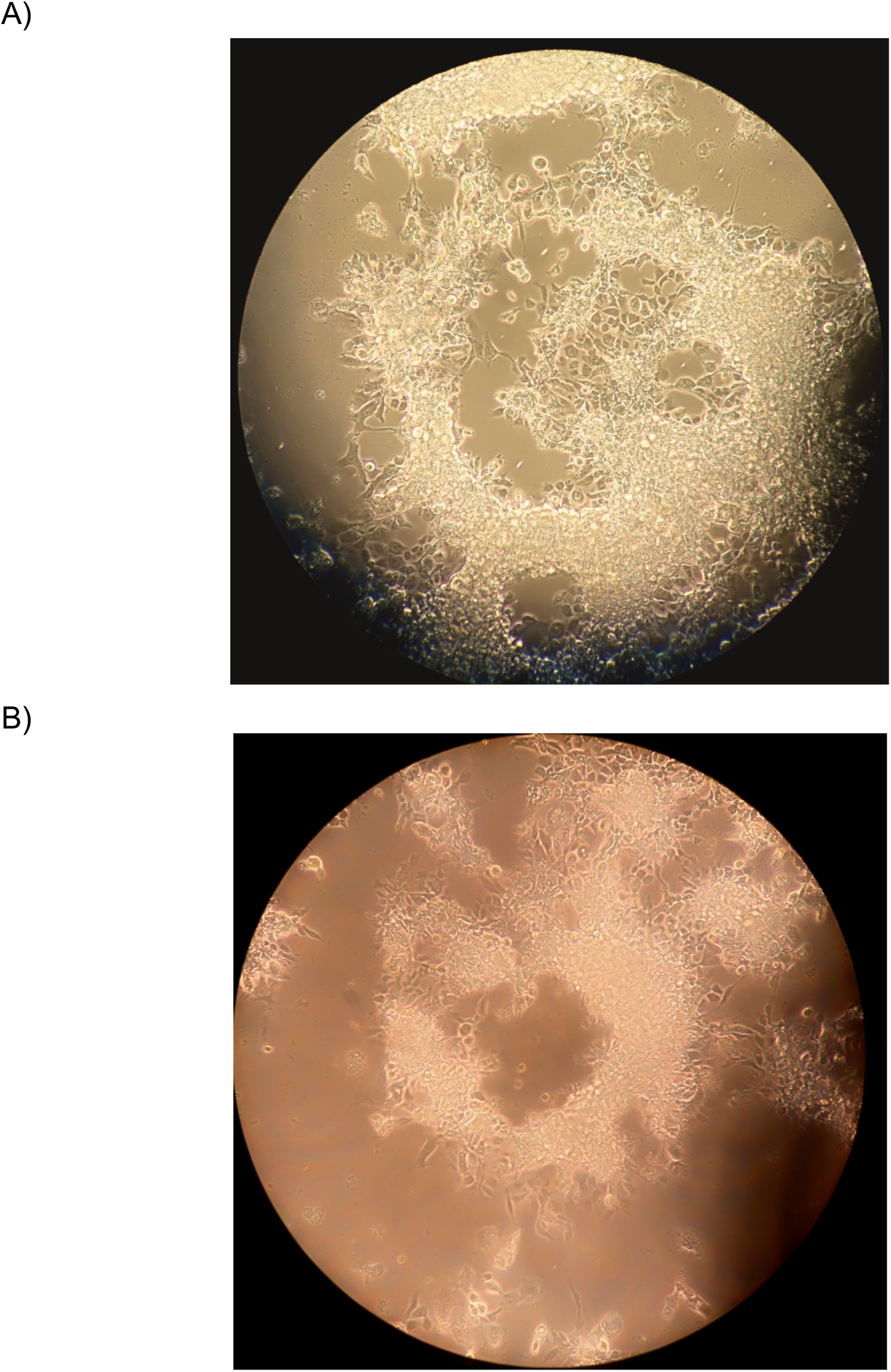
Aag2 culture cells 20X pictures. A) control, B) six hours post zymosan treatment.

## Notes

### Competing Interest Statement

The authors have declared no competing interest.

