## Supplementary Material for "*Aedes aegypti* Aag-2 culture cells enter endoreplication process upon pathogen challenge"

### *Drosophila melanogaster* hindsight protein orthologue corrected sequence in mosquito vector *Aedes aegypti*

<u>Introduction</u>

La endoreplicación en células epiteliales de intestino de mosquitos ha sido descrita como un probable mecanismo de respuesta inmune contra parásitos y virus.

Una de las cascadas de señalización posiblemente involucradas en la activación de síntesis de DNA genómico es la vía Delta-Notch.

Uno de los principales mediadores es el factor transcripcional *Hindsight* (molécula ortóloga de *Drosophila melanogaster*), cuya secuencia de RNA ha sido parcialmente descrita por herramientas bioinformáticas.

La proteína no ha sido descrita en *Aedes aegypti,* en la literatura de nuestro grupo se realizó una traducción reversa *in silico* para obtener la secuencia genómica más probable que codifica el mensajero de la proteína *hnt (Hindsight).*

El uso de la herramienta Exonerate permitió obtener una secuencia aminoacídica putativa de *hnt* (Figura Suplementaria A), mediante el mapeo de los fragmentos de DNA se obtuvo una región del supercontig 1.787 AaegL v.3.2.

Después se realizó la traducción de esta región no codificante con herramientas de predicción de exones, se obtuvieron hasta cinco secuencias distintas, además de la secuencia de *hnt de D. mel* (Figura Suplementaria B).

La región no codificante fue usada para traducir en los seis marcos de lectura para determinar el origen del RNA mensajero, el alineamiento de las proteínas putativas sugirió dominios conservados entre las secuencias (Figuras Suplementarias C, D, E, F y G).

Esto permitió identificar el marco de lectura y la secuencia completa de la proteína *hnt* en la región de DNA (Figura Suplementaria 1 y 2). La búsqueda de la proteína *hnt* en las bases de datos actualizadas en las publicaciones de referencia no arrojó ningún resultado.

A mediados de 2018, la disposición de una nueva base de datos mediante el trabajo de Matthew y colaboradores, permitió volver a realizar el alineamiento de la proteína putativa *Hindsight* mediante la metodología descrita anteriormente a la versión genómica AaegL5.2, con lo cual nos encontramos un porcentaje de identidad del 99.8762% al péptido AAEL020033 descrito por secuenciación del neurotranscriptoma del mosquito, pero sin función conocida a la fecha (Matthews et al. 2016, 2018).

Clustal Omega (Madeira et al. 2019), MView for spot differences in domains (Brown et al. 1998).

Usamos un anticuerpo dirigido contra la proteína *hnt* (DSHB, 1G9, AB_528278), usando como testigo intestinos de *Drosophila melanogaster,* no se observó señal en intestinos de *Aedes* *aegypti* con diferentes tratamientos (testigo solo con sacarosa al 10%, alimentados con sangre, infectados con DENV-2, *priming* con DENV-2).
